# Deep learning-based automated lesion segmentation on mouse stroke magnetic resonance images

**DOI:** 10.1101/2022.08.09.503140

**Authors:** Jeehye An, Leo Wendt, Georg Wiese, Tom Herold, Norman Rzepka, Susanne Mueller, Stefan Paul Koch, Christian J. Hoffmann, Christoph Harms, Philipp Boehm-Sturm

## Abstract

Magnetic resonance imaging (MRI) is widely used for ischemic stroke lesion detection in mice. A challenge is that lesion segmentation often relies on manual tracing by trained experts, which is labor-intensive, time-consuming, and prone to inter- and intra-rater variability. Here, we present a fully automated ischemic stroke lesion segmentation method for mouse T2-weighted MRI data. As an end-to-end deep learning approach, the automated lesion segmentation requires very little preprocessing and works directly on the raw MRI scans. We randomly split a large dataset of 382 MRI scans into a subset (n = 293) to train the automated lesion segmentation and a subset (n = 89) to evaluate its performance. We compared Dice coefficients and accuracy of lesion volume against manual segmentation, as well as its performance on an independent dataset from an open repository with different imaging characteristics. The automated lesion segmentation produced segmentation masks with a smooth, compact, and realistic appearance that are in high agreement with manual segmentation.

## Introduction

In ischemic stroke lesion detection, magnetic resonance imaging (MRI) is frequently used in both clinical and pre-clinical research. T2-weighted (T2w) MR images can be used to detect vasogenic edema and are often used as a proxy of the final infarct. In the hours after an ischemic stroke, the blood-brain barrier is disrupted and leads to fluid building up in the surrounding tissue. The resulting extracellular water increases T2 relaxation time, leading to increased signal in the stroke area. Ischemic brain tissue damage can be observed from T2w MR images as early as 3.5 hours after the onset of stroke and its volume continues to increase up to 24 hours after stroke^1^. T2w MR images acquired in this phase have become an important outcome parameter in rodent stroke research.

A challenge in stroke MRI research is that lesion segmentation often relies on manual tracing by trained experts. Manual segmentation is labor-intensive, time-consuming, and prone to inter- and intra-rater variability^2–4^. This challenge becomes especially problematic in pre-clinical research, which may involve large numbers of MR scans or multi-center datasets. Automated lesion segmentation is therefore crucial for both efficiency and reliability. Furthermore, automated lesion segmentation is essential for developing a fully automated processing of rodent stroke MR images. Atlas registration in the presence of a stroke lesion is challenging, and we have previously shown that this can only be effectively done if an algorithm is informed with a lesion mask^5^.

Several automated lesion segmentation algorithms have been developed for human data^6, 7^. With regards to animal data, there are only a few (semi-) automated methods. Jacobs et al.^8^ developed an unsupervised lesion segmentation model for rat data with an iterative self-organizing data analysis technique (ISODATA), a K-means derived clustering technique. A multiparametric ISODATA was applied to MR data to characterize ischemic lesion tissue in rats. Ghosh et al.^9, 10^ developed an automated ischemic lesion segmentation algorithm for neonatal rat brains using a hierarchical recursive region splitting (HRS) approach. Mulder et al.^2^ developed a level-set-based lesion segmentation algorithm for mouse data that requires one T2 MR sequence as input. Castaneda-Vega et al.^11^ trained a random forest classifier for ischemic stroke lesion segmentation in rat brains. However, most of these methods require some degree of manual intervention, require different parameters for each dataset for optimal performance, and are often dependent on the performance of other steps, such as preprocessing. Valverde et al.^4^ presented RatLesNetv2, a Convolutional neural network (CNN)-based Rodent Brain Lesion Segmentation algorithm for T2w MR images. However, RatLesNetv2 requires users to train their own model using their data.

In this exploratory study, we developed a fully automated ischemic stroke lesion segmentation method for mouse T2w MR images. The automated lesion segmentation builds on top of U-Net^12^, a convolutional neural network commonly used in biomedical image segmentation. We trained and evaluated the automated segmentation method on a large dataset of 382 T2w mouse MR images. Our approach produced realistic segmentations in high agreement with manually segmented lesion masks. The dataset used in this study is openly available on zenodo (https://doi.org/10.5281/zenodo.6379878) and presents one of the largest public T2w mouse stroke lesion repositories to date. This study thus contributes to making preclinical stroke lesion segmentation on MR images more standardized across studies and labs.

## Results

The resulting segmentation masks accurately detected lesions and produced a smooth, human annotator-like appearance. Incidence maps of manually segmented (median volume 26.73 mm^3^, IQR [7.65-48.27]) and automated (median volume 23.48 mm^3^, IQR [7.94-47.27) lesion masks are shown in Figure 2a. We also calculated false positives and false negatives across all evaluation datasets by subtracting manual lesion masks from automated lesion masks, shown in Figure 2b. Qualitative inspection of the spatial distribution showed the elevated incidence of false positives at the borders between cortical and subcortical structures and more homogenous distribution of false negatives across the lesion territory (Figure 2b).

**Figure 1.**
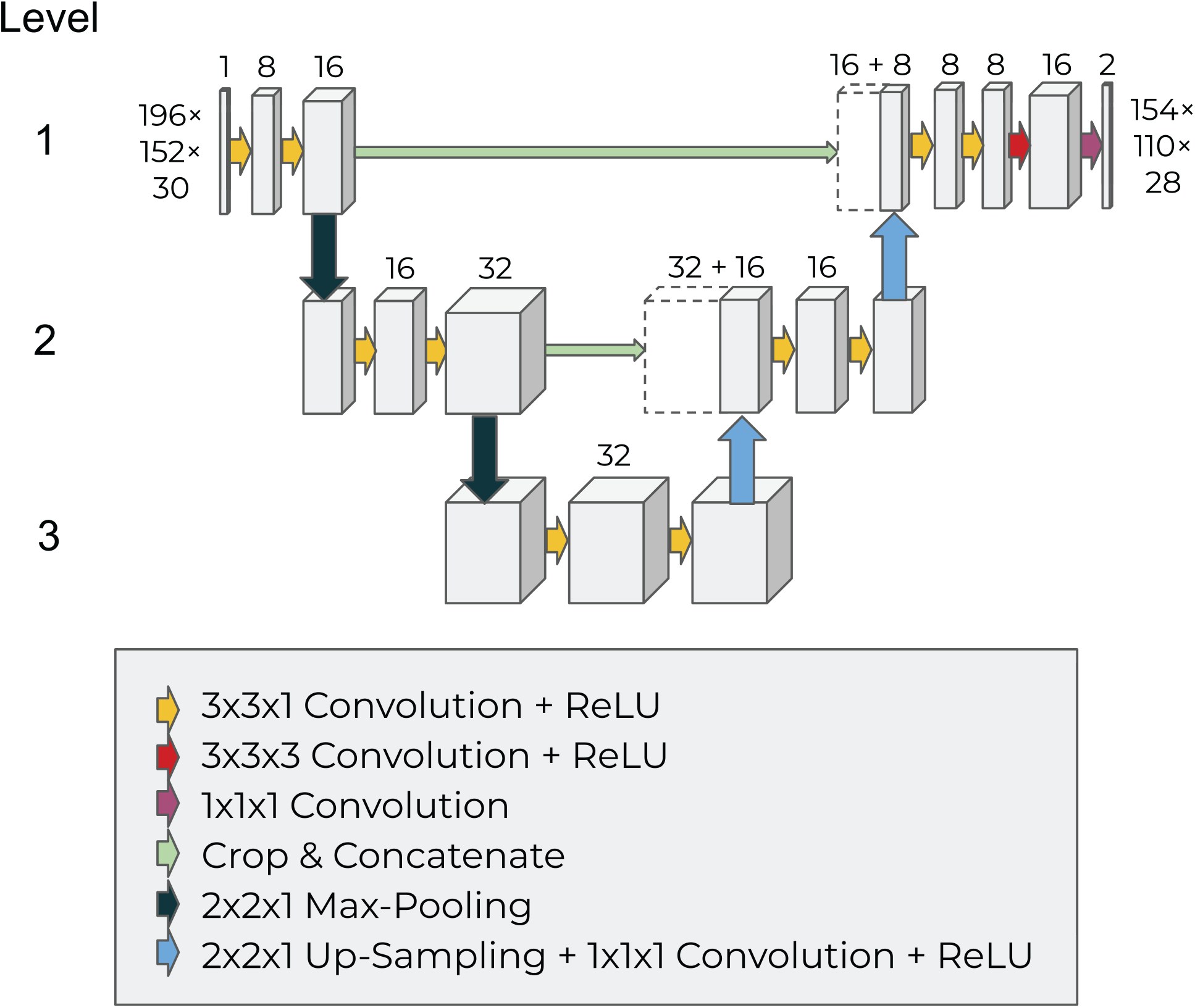
Network Architecture and processing pipeline of the automated method.

**Figure 2.**
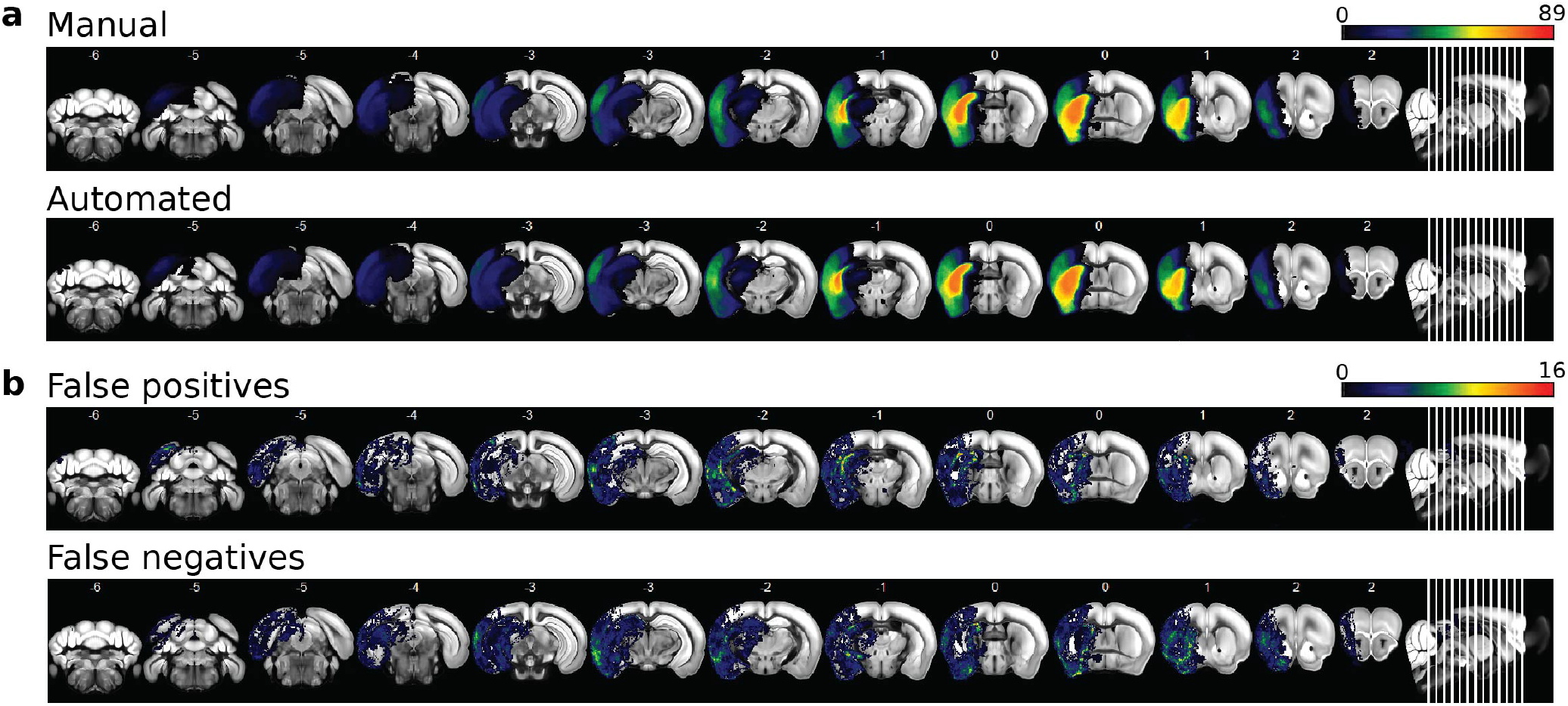
Incidence maps of (a) manual and automated lesion masks, and (b) false positives and false negatives across all evaluation datasets (n = 89).

The evaluation set included 12 scans without any lesions. Of those, the automated lesion segmentation detected small lesion masks in 10 datasets, with a median lesion volume of 0.34 mm^3^, IQR [0.19-1.45].

Figure 3a shows the Spearman correlation between manual and automated lesion mask volumes. The plot shows a very high level of correlation, with *ρ* = .98, n = 89, P = 4.81×*e*^−61^. The corresponding Bland-Altman plot (Figure 3b) indicates very high agreement between the two methods. No large systematic error was found but the automated algorithm tended to underestimate lesion volume for a few of the smaller lesions. Overall, we report a median Dice score of 0.92, IQR [0.86-0.96], a median sensitivity of 0.93, IQR [0.85-0.97], a median specificity of 1.00, IQR [1.00-1.00], and a median precision of 0.93, IQR [0.87-0.97]. In addition, we had an average Dice score of 0.89 ± 0.12.

**Figure 3.**
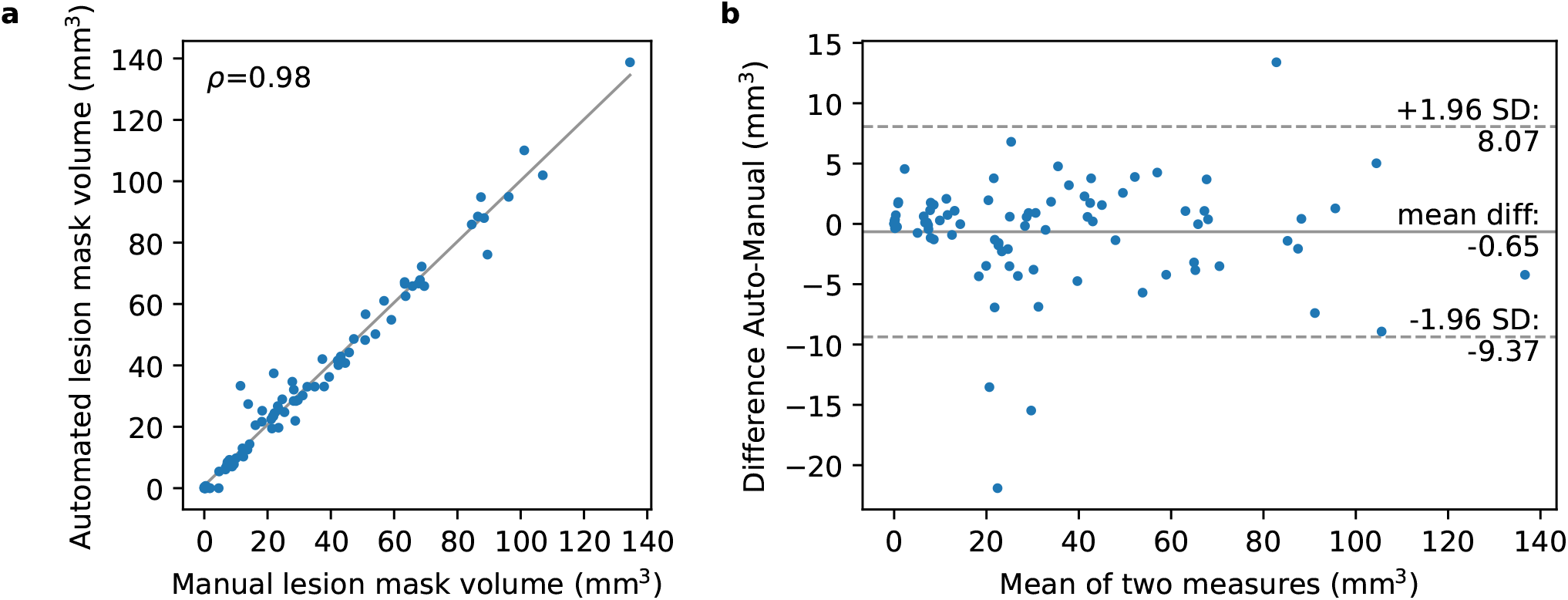
Relationship between the lesion mask volumes of the two methods (a) Spearman correlation and (B) Bland-Altman plot comparing agreement between the manual and automated lesion mask volumes.

To test the generalizability of our automated lesion segmentation method on other datasets, we evaluated our approach on an independent dataset^13^ and compared it with the performance of their method (see Methods section for more details). The results are shown in Table 1. Overall, the automated segmentation was able to segment the lesions reliably and with high accuracy. Our lesion masks tended to be smaller and more compact than lesion masks from their method, as shown in Figure 4. We had an average Dice score of 0.76 compared to 0.86 of Mulder et al.^2^.

**Table 1.** Comparison of performances of automated lesion segmentation algorithm by Mulder et al.^13^ vs. algorithm of the present study on open data repository of Mulder et al.^13^ Observer 1 vs. observer 2 refer to lesion masks produced by each of the two manual tracers from their study. The mean dice coefficient across animals within each group and the average of the mean dice coefficient across three groups are shown. For the present study, errors refer to standard deviation. For Mulder et al.,^2^ the type of error was not reported. We additionally report the range of dice scores with parenthesis.

| Age (months) | Time between infarct induction and MRI | Mulder et al. method |  | Present study |  |
| --- | --- | --- | --- | --- | --- |
|  |  | Observer 1 | Observer 2 | Observer 1 | Observer 2 |
| 3 | 18h (n=6) | 0.88 ± 0.03 | 0.89 ± 0.03 | 0.789 ± 0.05<br>(0.74 - 0.87) | 0.81 ± 0.05<br>(0.77 - 0.90) |
|  | 4d (n=3) | 0.85 ± 0.01 | 0.87 ± 0.01 | 0.68 ± 0.034<br>(0.63 - 0.72) | 0.712 ± 0.03<br>(0.69 - 0.75) |
| 1, 12 | 48h (n=10) | 0.86 ± 0.07 | 0.85 ± 0.07 | 0.80 ± 0.06<br>(0.66 - 0.87) | 0.76 ± 0.08<br>(0.64 - 0.87) |
| Average |  | 0.86 | 0.87 | 0.76 | 0.76 |

**Figure 4.**
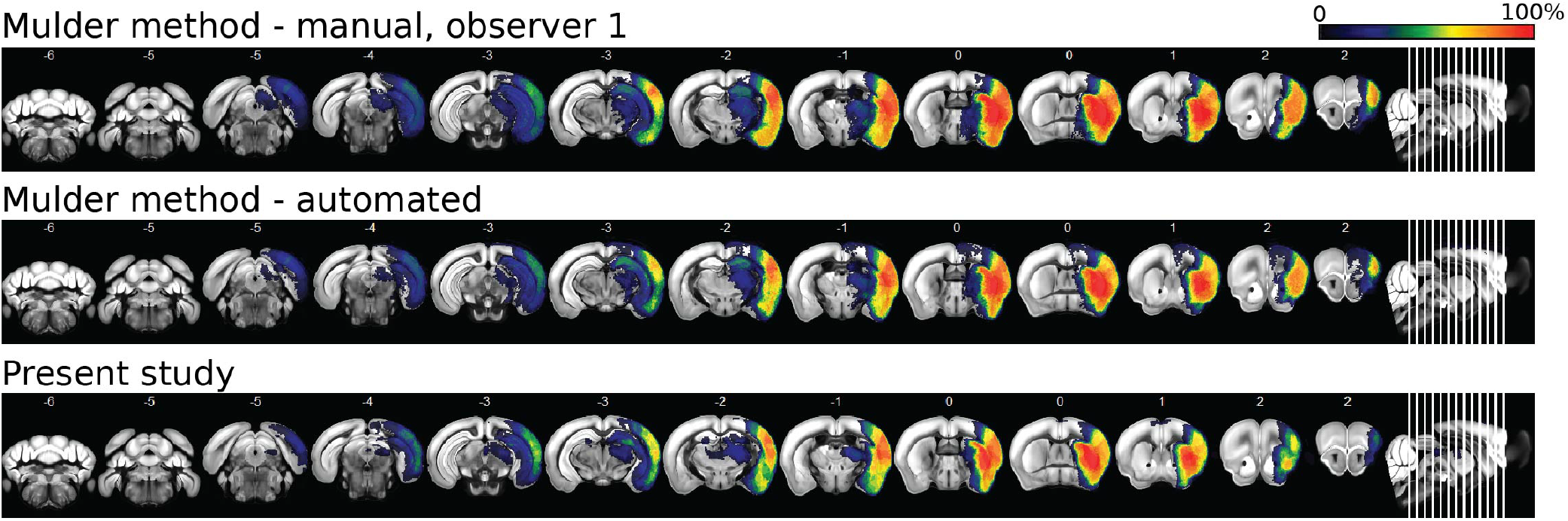
Incidence maps of automated lesion masks generated by Mulder et al.^2^ and by the present study.

## Discussion

We have presented an automated algorithm for ischemic lesion segmentation from mouse T2w MR images. Our reported median Dice score of 0.92 shows that the automated lesion segmentation produces realistic segmentations that very closely resemble human annotations. In contrast, previous studies showed that annotations made by two human annotators can have a Dice score as low as 0.73 or 0.79^2, 4^.

The convolutional neural network architecture used in our approach is based on a variation of the 3D U-Net^12^ design, a highly successful segmentation model for bioimaging tasks. As an end-to-end deep learning approach, the automated lesion segmentation requires very little preprocessing and works directly on the raw MRI scans. Unlike some of the previous automated lesion segmentation approaches^2, 9^, the present study is fully automated and it is not dependent on other preprocessing steps, such as registration or skull stripping. This is crucial in achieving a fully automated pipeline for processing rodent stroke MR images, especially since we have previously shown that such preprocessing steps are prone to error in presence of a stroke lesion^5^.

Our approach is similar in design to the recently published RatLesNetv2-architecture^4^ developed for rat data. However, we used a much larger training set and incorporated 3D information across the z-axis into our predictions. They report average dice scores (inter-rater: 0.73 ±0.12, RatLesNetv2: 0.81 ± 0.16) on their evaluation set that are lower than our best-performing evaluation score (average dice score of 0.89±0.12).

We further tested the automated lesion segmentation on the dataset provided by Mulder et al.^13^ and tested generalizability across scans from a different center with different imaging characteristics (lower resolution; T2 maps instead of T2w images; different scan times after infarct induction). Our Dice score on Mulder et al.’s data^13^ fell short of their performance (Table 1). However, their ground-truth annotations contain holes, rough and fringed edges, and spots of isolated voxels since a fixed intensity threshold based on contralateral mean and standard deviation was used without further postprocessing. These differ considerably from ground-truth labels in our dataset, which mostly contain compact lesion masks with smooth edges. We defend the use of compact lesion masks based on the following two arguments. First, consistent with literature^4, 14^, we argue compact lesion masks are more realistic and comparable to manual annotations since hypoperfusion in large vessel stroke varies smoothly in space and not at the order of single voxels. Second, our manual lesion masks were obtained using a semi-automated manner using a thresholding function and the lesion masks obtained using the same procedure have been validated by comparison to histological data using TTC staining in a previous study^5^, where we report an excellent correlation. Given that TTC shows very little variation on the scale of a single voxel, this further supports our assumption of compact masks being consistent with the ground truth. We would also like to note that the dice score reported by Mulder et al.^2^ is obtained from the same data that was also used to train and optimize the model. Meanwhile, our model shows reliable performance despite being trained using a different dataset.

The comparison between our automated method and other previously published methods is limited to the application of our algorithm to the data published by Mulder et al.^13^. To our knowledge, Mulder et al.^2^ is the only end-to-end approach developed for mice data thus far. We could not test our data using Mulder et al.^2^’s method since their algorithm only takes T2 maps as input which were unavailable for our data. This points to another advantage of our automated method, which is that it can take any image in NIfTI format as input, including T2w images and T2 maps, on which the stroke appears hyperintense.

T2w images acquired at this time point need to be corrected for the effect of brain edema since swelling due to post-stroke edema leads to a significant overestimation of lesion volume compared to histology^11, 15^. A study showed that hyperintensity in T2w at approximately 7 h after stroke represents the final lesion volume and that the subsequent increase in the signal would mainly reflect swelling^1^. The correction could be performed using the atlas registration approach by Koch et al.^5^ or the model by Gerriets et al.^15^. With this consideration in mind, T2w images are the most commonly used imaging technique to estimate the final infarct in animal studies of experimental stroke^16^ and and serve as a useful measure for correlation of lesion site with other imaging biomarkers.

Diffusion Weighted Imaging (DWI), in addition to T2w images, is also routinely applied for ischemic stroke segmentation. Whereas T2w images are sensitive to vasogenic edema, apparent diffusion coefficient (ADC) maps derived from DWI indicate cytotoxic edema, i.e. water diffusion restrictions due to cellular swelling. A reduction in ADC is observed in the first few hours after stroke induction and it is shown to be a strong predictor of final lesion volume^17, 18^. ADC and T2w data measure partly overlapping and distinct microstructural lesion traits, and lesion segmentation from ADC and T2w show different patterns at 24 hours after stroke^8, 11^. In addition, the combined evaluation of ADC and T2w is shown to lead to a more accurate prediction of the final stroke volume^8, 11^.

To test whether the method could also generalize across different MR contrasts, we ran the automated method on a preliminary dataset of 5 DWI acquired after permanent distal MCAo in the same mouse model. Here, the automated method showed poor performance, with an average dice score of 0.24. Thus, a limitation of our method is that, although we show that it could be generalized across data sources and small changes in surgery, it may not generalize well across different MR contrasts and surgery procedures.

Another limitation of our approach is that it is only end-to-end if B1+/- is homogenous. Intensity variation due to B1+/- and bias field issues are common in high-field MR images acquired with a preclinical scanner. With the presence of B1+/- inhomogeneity, an extra preprocessing step involving B1 correction, e.g. using ANTs N4 Bias Field correction^19^ or SPM’s bias field correction using simultaneous segmentation/registration procedure^20^, is necessary.

Finally, a limitation is that we did not have ground truth based on histology for this study, and the manual segmentations from MRI served as the "ground truth" for model training. However, we would like to stress again that the manual segmentations obtained through the same method have been validated against histology data in a previous study^5^, where we report high correlation between the lesion volumes.

To conclude, a fully automated end-to-end deep learning approach to segment stroke lesions in preclinical studies has been trained on one of the largest datasets of T2w MR images acquired in subacute murine stroke to date. The method performed well compared to manual segmentation and will help to standardize MRI-based lesion volumetry between studies and labs. Furthermore, it can now be used to inform other algorithms needed for rodent MR image preprocessing that previously relied on manual interaction.

## Methods

### Animals

We retrospectively reanalyzed male C57/BL6 J mice (Janvier, Germany) from previous stroke studies, most of which are unpublished. Animals underwent middle cerebral artery occlusion (MCAo, n = 332), or a sham procedure (n = 50) at the age of 10–12 weeks. Experimenters were blinded to the condition of the animals. Mice were housed in a temperature (22±2°C), humidity (55±10%), and light (12/12-h light/dark cycle) controlled environment. All surgeries were performed under 1.5%–2% isoflurane in a 70% nitrous oxide and 30% oxygen mixture. During surgery, core body temperature was maintained at 37±0.2°C with an automated rectal probe and heat blanket. After surgery, a topical application of 1% bupivaccain gel was applied on surgical wounds. All experiments were approved by the Landesamt für Gesundheit und Soziales (registration numbers G0197/12, G0005/16, G0057/16, G0119/16, G00254/16, G0157/17, and G0343/17) and were performed in accordance with the German Animal Welfare Act and the ARRIVE^21^.

### MCAo

After anesthesia induction, a midline neck incision was made to expose the left carotid artery. A filament (190 μm diameter, Doccol, Sharon, MA/USA) was introduced into the external carotid artery (ECA) and advanced to the origin of the left MCA via the internal carotid artery. After 45 min, the filament was withdrawn. For sham animals, the filament was advanced to the MCA and withdrawn immediately. Both the common carotid artery and the ECA were ligated and remained occluded during reperfusion^22^. All MCAo were performed by two experienced surgeons.

### MRI measurements

Mice were anesthetized with isoflurane (2-2.5% at initiation and 1-2% during the scanning period) in a 70:30 nitrous oxide:oxygen mixture. Respiration rate was monitored with a pressure-sensitive pad using MRI compatible equipment (Small Animal Instruments, Inc., Stony Brook, NY, USA). T2w images were acquired 24 h post-stroke surgery on a 7 T MR scanner (Bruker, Ettlingen, Germany) and a 20 mm inner diameter transmit/receive quadrature volume coil (RAPID Biomedical, Rimpar, Germany). A 2D RARE sequence was used (repetition time/echo spacing/effective echo time = 4.2 s/12 ms/36 ms, RARE factor = 8, 32 consecutive 0.5 mm thick slices, FOV = (25.6 mm^2^), image matrix = 256 × 196 zerofilled to 256 × 256, 4 averages, scan time 6:43 min).

### Manual lesion segmentation

The manual lesion masks were created in a semi-automated manner by tracing hyperintense lesions using a thresholding function in ANALYZE 10.0 software (AnalyzeDirect, Overland Park, KS, USA). The lesion masks were of the same dimensionality as the T2w images and were exported in NIfTI format. The manual segmentation was performed by an experienced researcher with over 20 years of experience and whose tracings were validated against histology in previous studies^5^. Each manual segmentation took around 5 minutes.

### Independent dataset

An independent dataset from a previous study^13^ was used to further evaluate the automated lesion segmentation. This independent dataset is different from that of the present study in that it is of lower resolution (128×128 and 196×196, instead of 256×256), contains quantitative T2 maps (instead of T2w images), and had different scan times after the infarct induction. The authors provide two manual annotations from independent observers, which were segmented by applying various thresholding operations in ImageJ in a semi-automated manner. Some of the low signal-to-noise ratio data in the dataset, therefore, contained very fragmented lesion masks with small holes (order of one or few voxels). We evaluated the automated lesion segmentation on a subset of 19 higher-quality datasets (Cologne-Set 1, Cologne-Set 2), which we found to have lesion masks that were less fragmented, and thus more realistic, than other datasets.

### Automated lesion segmentation

#### Network architecture

In order to automatically segment the lesion, a variant of the popular U-Net was trained^12^. The architecture follows an encoder-decoder design but adds "skip connections" between intermediate layers of the same spatial resolution.

As shown in Figure 1, two max-pooling layers were applied, allowing the network to analyze the image at three distinct spatial resolutions. The upsampling was implemented as nearest-neighbor upsampling, followed by a convolutional layer with kernel size 1×1×1 and ReLU activation. Between consecutive downsampling or upsampling steps, two convolutions with kernel size 3×3×1 and ReLU activation were applied. At last, a 3×3×3 convolution with ReLU activation was applied, followed by a 1×1×1 convolution with a softmax activation, the output of which is interpreted as the background and lesion probabilities. All in all, the network maps a 196×152×30 image tensor to a 154×110×28×2 tensor of probabilities. The receptive field of the network is 43 × 43 × 3 voxels which equals 4.3 × 4.3 × 1.5 mm^3^.

The number of feature maps was set to 8, 16, and 32, depending on the level of the U-Net. The number of feature maps of the next level was already used on the last convolution before the down-sampling^23^. Batch normalization layers were purposefully excluded because we experienced issues in low-contrast parts of the dataset when batch normalization was included. The network had 42,442 trainable parameters.

#### Model training

The boxes were split randomly into a training (n = 293) and validation set (n = 89). The (0, 255) interval of intensity values was scaled to the (−1, 1) interval by transforming them via I := (I - 127.5) / 127. As a data augmentation, voxel-wise Gaussian noise was added to the training samples with a mean of 0 and a standard deviation of 0.45.

The voxel-wise loss was computed as the categorical cross-entropy between the predicted probabilities and the one-hot encoded ground truth class. The training loss was defined as the total categorical cross-entropy across all voxels in a batch of eight training examples. Voxels were weighted inversely proportional to the frequency of the respective class.

All kernels were initialized according to Glorot and Bengio^24^. Biases were initially set to zero. The network was trained to minimize the training loss using the Adam optimizer^25^ with a learning rate of 1e-4 and a batch size of 8.

#### Implementation

The automated lesion segmentation method was implemented in TensorFlow and Python using the Voxelytics framework (https://voxelytics.com; scalable minds, Potsdam, Germany). Training and predictions were executed on a system with an NVidia Geforce GTX1080, an Intel Core i7-6700 CPU @ 3.40GHz, and 64GB RAM.

### Qualitative comparison and Statistical Analysis

T2w images and corresponding lesion masks were registered to the Allen mouse brain atlas using ANTx2 as described previously (https://github.com/ChariteExpMri/antx2)^5^. Incidence maps were generated by displaying voxel-wise mean of lesion masks across animals (expressed in percent) in atlas space. For each animal, a map of false positives was generated by voxel-wise subtraction of the manually generated lesion mask from the automatically generated lesion mask in atlas space followed by removal of all values < 1. Incidence maps of false positives were then generated by voxel-wise mean of false positive maps across animals (expressed in percent). For incidence maps of false negatives, the same procedure was performed except the automatically generated mask was subtracted from the manually generated mask.

We evaluated the automated lesion segmentation on a dedicated evaluation set consisting of 89 datasets. We used several metrics for assessment. First, we used total lesion volume, which is one of the most important readout parameters in preclinical stroke studies. Since the lesion volumes are not normally distributed, the manual and automated lesion volumes were compared using Spearman’s rank correlation and Bland-Altman analyses. Correlation and Bland-Altman analyses and visualization were performed using Python (version 3.8.8). Secondly, we used the Dice coefficient^26^, which measures the overlap volume between two regions and is popularly used for performance evaluation in the field of image segmentation. In addition, we report sensitivity, specificity, and precision rate. Sensitivity evaluates the capability to correctly detect true positives, whereas sensitivity focuses on the correct detection of true negative voxels. Precision is a measure of reliability that evaluates the fraction of predicted positives that is actually positive. As these measures are not normally distributed we report them with median and interquartile range (IQR). For comparison with previous studies that report only the average dice score, we also report the average dice score ± standard deviation.

## Data Availability

The datasets used for this study can be found in the zenodo repository, https://doi.org/10.5281/zenodo.6379878.

## Acknowledgements

Excellent technical assistance was provided by Janet Lips, Larissa Mosch, Monika Dopatka and Marco Foddis. Ulrich Dirnagl initiated the collaboration between Charité-Universitätsmedizin Berlin and scalable minds GmbH.

This study was supported by the Brandenburg Ministry of Economic Affairs, Labour and Energy (Brandenburgischer Innovationsgutschein) and the European Regional Development Fund. Funding to JA, SM, SPK and PBS was provided by the German Federal Ministry of Education and Research under the ERA-NET NEURON scheme (BMBF, 01EW1811), the German Research Foundation (DFG, Project HO5277/3-1, project number 417284923 to CJH, HA5741/5-1, project number 417284923 to CH, Project-ID 424778381 –TRR 295 to CH, BO4484/2-1, project number 428869206 to PBS, the Open Access Publication Fund of Charité - Universitätsmedizin Berlin and EXC-2049 – 390688087 NeuroCure) and by a grant from the Fondation Leducq (17 CVD 03) to CH. Retrospective analyses of previously acquired animal data were supported by Charité 3R| Replace - Reduce – Refine.

## Author contributions statement

JA, TH, NR, PBS. designed the study, CH, CJH and SM provided and annotated data, JA, TH, NR, SPK, PBS preprocessed and organized the data, TH, NR, GW, LW wrote the software and analyzed data, JA and SPK performed the statistical analyses, JA, GW, TH and PBS wrote the manuscript, all authors commented to the manuscript at all stages.

